# Environmentally-Friendly Workflow Based on Supercritical Fluid Chromatography and Tandem Mass Spectrometry Molecular Networking For the Discovery of Potent Anti-Viral Leads From Plants

**DOI:** 10.1101/106153

**Authors:** Louis-Félix Nothias, Stéphanie Boutet-Mercey, Xavier Cachet, Erick De La Torre, Laurent Laboureur, Jean-François Gallard, Pascal Retailleau, Alain Brunelle, Pieter C. Dorrestein, Jean Costa, Luis M. Bedoya, Fanny Roussi, Pieter Leyssen, José Alcami, Julien Paolini, Marc Litaudon, David Touboul

## Abstract

A supercritical fluid chromatography-based targeted purification workflow using tandem mass spectrometry and molecular networking was developed to analyze, annotate and isolate secondary metabolites from complex mixture. This approach was applied for targeted isolation of new antiviral diterpene esters from *Euphorbia semiperfoliata* whole plant extract. The analysis of bioactive fractions revealed that unknown diterpene esters, including jatrophane esters and phorboids esters, were present in the samples. The purification procedure using semi-preparative-supercritical fluid chromatography led to the isolation and identification of two jatrophane esters (**13** and **14**) and four 4-deoxyphorbol esters (**15**-**18**). Compound **16** was found to display antiviral activity against chikungunya virus (EC_50_ = 0.45 *µ*M), while compound **15** was found to be a potent and selective inhibitor of HIV-1 replication in a recombinant virus assay (EC_50_ = 13 nM). This study showed that supercritical fluid chromatography-based workflow and molecular networking can facilitate and accelerate the discovery of bioactive small molecules by targeted molecules of interest, while minimizing the use of toxic solvents.

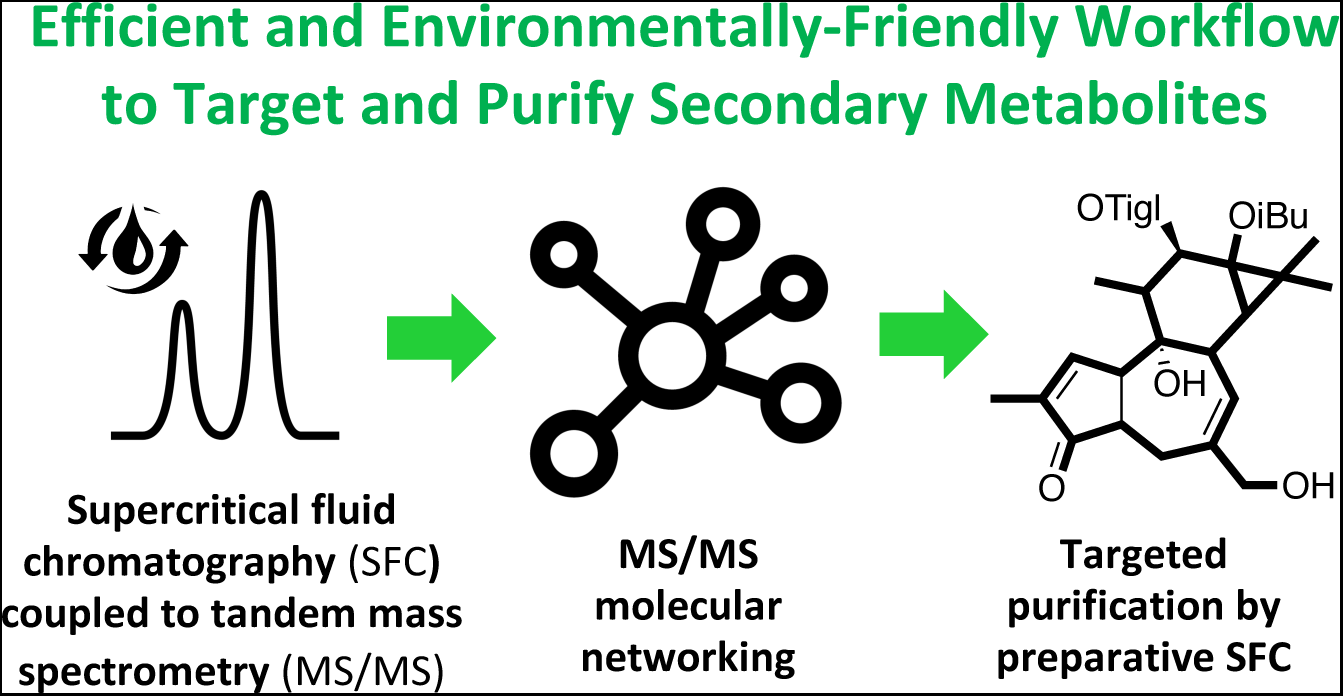

---

The secondary metabolites produced by various organisms such as plants, bacteria, fungi, and insects, represent a unique reservoir of chemical diversity, often endowed with potent biological properties.^1^ If metabolomics can help deciphering structural diversity from a complex mixture,^2,3^ the discovery of new bioactive molecules is still a very challenging task. The isolation of bioactive natural products is generally achieved by a bioassay-guided purification procedure.^1^ However, this procedure is not always successful, often leading to isolation of known molecules. The whole process is time-consuming, and generally uses significant amount of toxic organic solvents. Last decades, hyphenated analytical methods such as liquid chromatography coupled to mass spectrometry (LC-MS) or NMR (LC-NMR) have been used as dereplication tools to detect the presence of known bioactive molecules prior to start any isolation procedure.^3,4^ However, despite the development of efficient LC-MS-based method for generic and comprehensive profiling of secondary metabolites in natural extracts,^5^ the annotation of new and known analogues in a systematic and untargeted way remains a challenging task.

For these reasons, the development of more efficient approaches is highly required to discover new bioactive molecules. To overcome the current limitations of existing methodology, these new approaches should be able (i) to prioritize biomass selection by facilitating the annotation of analogues, and the detection of unique chemical structure, (ii) to accelerate isolation of bioactive compounds, (iii) to minimize potential degradation of bioactive compounds, and finally (iv) to minimize the use of toxic chemicals reducing the environmental impact of the overall process.

Supercritical CO_2_ (sCO_2_) is now widely used in industry allowing the extraction of natural products with a reduced environmental impact^6–9^ or the separation of chiral compounds at analytical or preparative scales.^10^ Over the last few Years, supercritical fluid chromatography (SFC) is gaining back of interest because manufacturers (Agilent Technologies, Waters, and Shimadzu, in particular) marketed user-friendly and robust analytical and preparative systems. Compared to LC, SFC offers a largest choice of stationary phases exhibiting different selectivity allowing better chromatographic resolution and shortened retention times due to the low viscosity and high diffusivity of sCO_2_. Moreover, up-scaling from analytical to (semi)preparative scale is achievable while preserving a good chromatographic resolution. Although terpenes constitute a large class of non-polar compounds, publications dealing with their separation and isolation process using sCO_2_ are scarce, except for monoterpenes.^11,12^

Like HPLC system,^13^ SFC-based system can be coupled with tandem mass spectrometers in order to provide structural information on biomolecules. While bioinformatics tools are now well implemented for metabolomics,^14^ or proteomics,^15^ only few of them can really empower the information covered in MS/MS data of natural compounds. Among them, MS/MS molecular networking (MS^2^MN) has emerged as an efficient tool to explore MS/MS data since its availability on Global Natural Product Social molecular networking web-platform (GNPS).^16–20^ MS^2^MN is a graph-based approach that relies on the assumption that structurally related molecules share similar fragment ions in their respective MS/MS spectra.^16,21^ Thus, MS^2^MN allows establishing spectral similarity relationships represented as a spectral networks, so-called molecular networks (MNs). The emergence of this approach unleashed the potential of untargeted MS/MS analysis by allowing the visualization of spectral groups in an untargeted way. Moreover, MNs can be seeded with MS/MS data of molecular standards used to seed the MNs and propagate annotation,^18^ leading to the dereplication of not only identical molecules but also analogues organized as molecular series.^17^ Recent studies successfully employed MS^2^MN annotation in LC-MS/MS-based metabolomics and natural product isolation,^20,17,18^ but none of them was based on SFC-MS/MS data.^22–25^

We will expose our efforts to develop an efficient and environmentally friendly SFC-MS^2^MN-based workflow for the discovery and isolation of new bioactive molecules from a complex natural extract. The study of *E. semiperfoliata* plant extract endowed with strong antiviral activity^26,27^ offers an interesting subject to optimize and evaluate our approach. In fact, several chromatographic fractions showing antiviral potency were not studied yet,^28^ while phorboids diterpene esters were reported from this species but never evaluated.^29^ Moreover, thanks to the described MS/MS fragmentation behavior of some jatrophane esters previously isolated from this species, this study also aimed at exploring how diterpene ester fragmentation pattern drive molecular network clustering.

## RESULTS AND DISCUSSION

### Development of a SFC-MS/MS Method for the Analysis of Diterpene Esters

In order to determine the best conditions of separation, various parameters, *i.e.* stationary phases, the mobile phase composition and flow-rate, back pressure and oven temperature, were optimized using a test-mixture of seven jatrophane esters standards (**3**-**8**, and **11** in Figure 1).^28^

**Figure 1.**
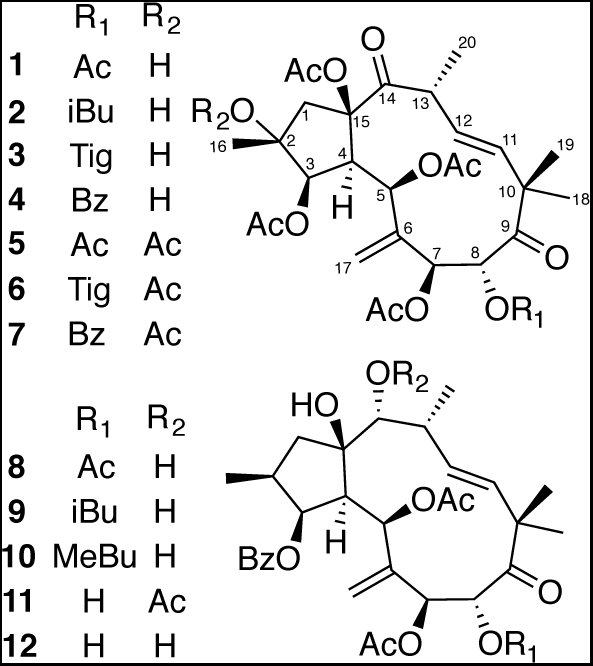
Jatrophane esters isolated from *Euphorbia semiperfoliata*, compounds **3-8** and **11** were used as reference standards in this investigation.

Figure 2 shows that very low selectivity was obtained using CN, PFP, 2-EP, 1-AA, 2-PIC, DEA and C18 stationary phases, while intermediate selectivity was obtained using Si and DPH using generic gradient from 5 % to 20 % of ethanol in 20 minutes. Zr-Carbon and Hypercarb^®^ columns demonstrated excellent and similar selectivity. These two stationary phases bear hydrophobic surfaces and highly delocalized π-electron system. Because this column gave better peak widths and symmetries, Hypercarb^®^ was selected for further optimization.

**Figure 2.**
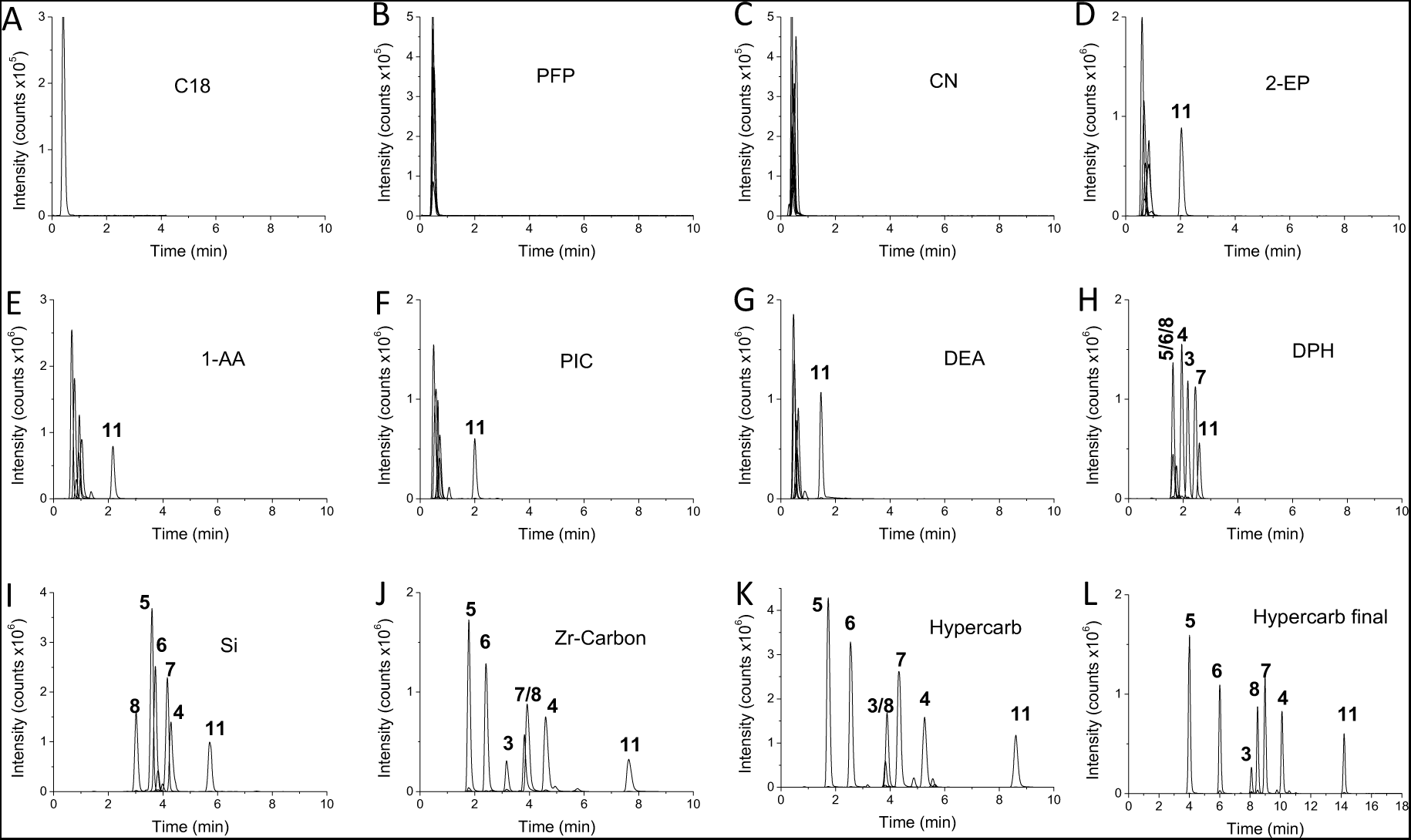
Extracted-ion chromatograms of jatrophane esters **3-8** and **11** obtained by SFC-MS using 11 different stationary phases (A-K), and the optimized method using Hypercarb column (L).

Optimal separation (Figure 2, thumbnail L) was obtained by decreasing the particle size down to 3 µm and adjusting the gradient of the mobile phase as described in the experimental section. The relative standard deviation on retention times over five successive injections was always below 1 % indicating that no alignment is necessary for data retreatment. Carry-over was estimated to be below 1 %.

### Molecular Networking with SFC-MS/MS data

The ethyl acetate extract of the whole plant of *E. semiperfoliata* showed significant antiviral activities against CHIKV and HIV replication, and some of the isolated jatrophane esters were shown to display selective activities.^28,32^ In the present study, MS^2^MN was used to explore the molecular content of bioactive fractions obtained previously from this extract.28,32 Analysis by SFC-MS/MS were performed on standard compounds andfractions, and data were further analyzed by on GNPS web platform.^19^

Fractions F7, F8 and F9 exhibiting potent antiviral activity against CHIKV virus replication (EC_50_ = 1.2, 9.4 and < 0.8 *µ*g.mL^-1^, respectively) were selected. The MS/MS data were acquired using a data-dependent acquisition method to collect collision-induced fragmentation spectra for the four most intense ions observed in the range *m*/z 300-800.19 MS/MS spectra of previously isolated compounds 1-12 were included in the dataset as references to annotate some clusters, and to optimize MS^2^MN parameters. The Δ
*m/z* between two nodes in the MN were annotated as molecular differences using a table of elemental compositions (elements C, H, O, N, and Na only),30 according to mass errors below 0.01 Da.

The use of default parameters of the GNPS workflow led to the generation of molecular network nodes that were found to be artefactual. Indeed, some consensus MS/MS spectra clusterered by MS-Cluster, were of chimeric nature. The chimeric MS/MS spectra are due to the clustering of MS/MS spectra produced by different, but related molecules, with a cosine score above 0.7, and precursor ions within the precursor ion mass tolerance (2.0 Da). After optimization of molecular networks parameters including the use of a *precursor ion mass tolerance* of 0.01 Da, and a *product ion tolerance* of 0.0075 Da, the number of these chimeric features could be drastically reduced.

Furthermore, to achieve a satisfactory clustering of diterpene esters using reference-standard compounds, molecular networks were generated using 12 minimum matched peaks, and by removing fragment ions below 10 counts from the MS/MS spectra.

The MS^2^MN of fractions F7-F9 are shown in Figure 3. Annotation of the known jatrophane esters in the MS^2^MN was allowed by spectral matching with the MS/MS spectra of reference-standards (**1**-**12**), and by comparison with retention time obtained in the same experimental conditions, allowing a level 1 identification based on Metabolomic Standards Initiatives.^31^ Moreover, annotation (level 2) was propagated through the network by inspection of the MS/MS spectra of nodes connected to reference standards. Indeed, it has been shown that groups A (blue color) and B (purple color) of jatrophane esters have different diterpene backbone fingerprint, with characteristic fragment ions at *m/z* 327, 309 and 299 (group A), and *m/z* 313, 295 and 285 (group B).32 Jatrophane esters of group A possess a ketocarbonyl at C-14 and an acyl group or an hydroxy group at C-2, while jatrophane esters of group B possess an acyl group or an hydroxy group at C-14, and no additional acyl group at C-2. The main differences observed in elemental composition between nodes annotated as analogues were 2H, O, CH_2_, C_2_H_2_O, suggesting structural variation of the functional groups, such as a double bond, hydroxylation, length of acyl chain, or additional acetylation, respectively.

**Figure 3.**
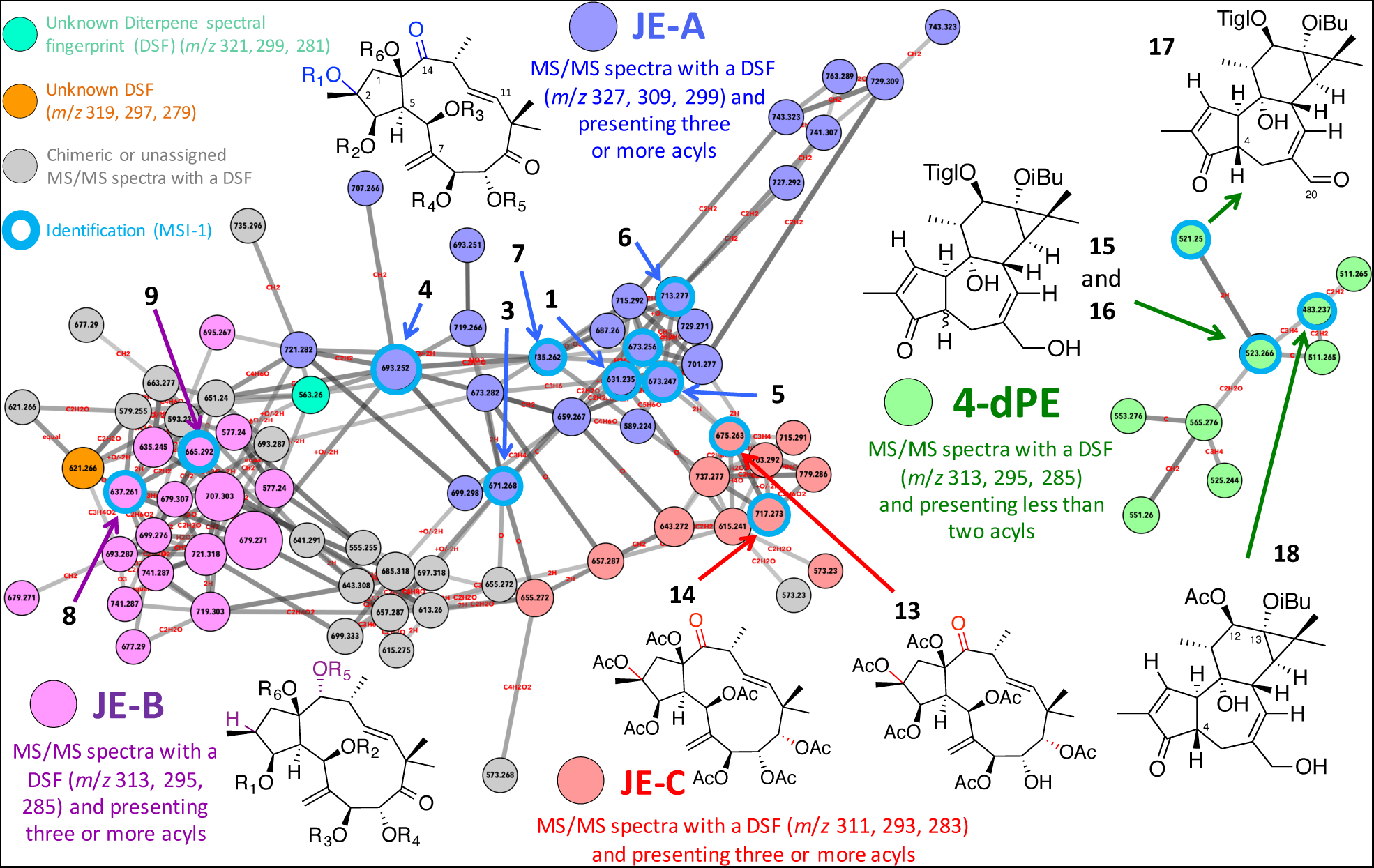
Annotation of the MS^2^MN of antiviral fractions F7-F9 derived from *E.semiperfoliata* extract showing jatrophane esters (JE) of group A (JE-A, **1**-**7**), and JE ofgroup B (JE-B, **8**-**9**), and their putative analogues. New compounds, jatrophane esters (JE) of group C (JE-C, **13**, **14**) and 4-deoxyphorbol esters (4-dPE, **15** -**18**), were targeted upstream during the initial interpretation of MS^2^MN of SFC-MS/MS analyze, and were further isolated by semiprep-SFC. MSI-1 and MSI-2: Metabolomic Standard Initiative identification level.

A third cluster (red color) showed MS/MS spectra evoking diterpene backbone fingerprint of phorbol esters (*m/z* 311, 293 and 283),^32,33^ but was interpreted as unknown jatrophane esters, because these spectra contained more than four neutral loss of acyls. Indeed, the maximum number of ester groups found in phorbol derivatives usually does not exceed three,^32^ Thus, these molecules were annotated as modified jatrophane esters of group C. Comprehensive inspection of each node in the main MN, revealed that multiple nodes (in grey) were a result of consensus MS/MS spectra produced during the MS-clustering process. These nodes are composed of nearly identical MS/MS spectra of related molecules possessing the same precursor ion, but different diterpene backbone fingerprint and retention time. The second MN (green color) contained MS/MS spectra reminiscent of compounds with the same apparent diterpene backbone fingerprint as jatrophane esters of group B (*m/z* 313, 295 and 285)^32,33^ but the MS/MS spectra of these compounds differed from the jatrophane esters of group B by the presence of only two or less loss of acyl-groups. Moreover, the MS/MS spectra of the ion at *m/z* 523.266 was consistent with a structure of 4-deoxyphorbol ester, which was previously isolated by Appendino et al.^29^ (Figure 4). This MN was tentatively annotated as 4-deoxyphorbol esters according to these observations. Although the clustering of diterpene esters in MS/MS molecular networks appeared to be driven by (i) the type of diterpene esters, and (ii) their functionalization, this approach is not sufficient to establish the molecular structure of these annotated compounds. We first aimed at selecting the best fraction for a targeted SFC-MS-based purification procedure.

**Figure 4.**
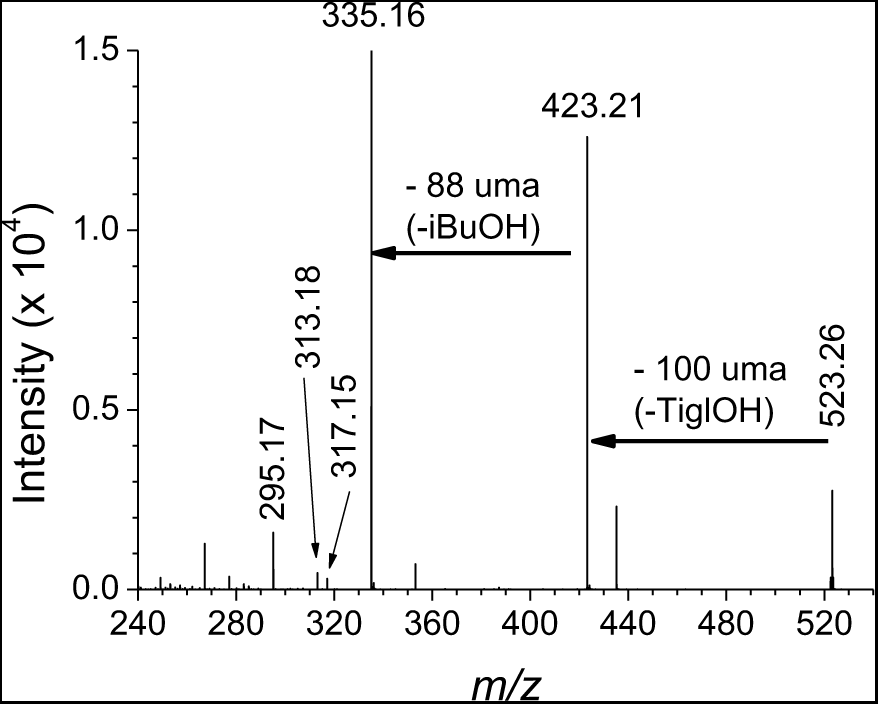
MS/MS spectrum of the precursor ion at *m/z* 523.27 (compound **16**),putatively annotated as 4-deoxyphorbol diester with tiglyloxy and isobutyryloxy moieties.

In order to give information on the relative distribution of nodes between the three fractions, the data-dependent MS/MS acquisition method were achieved with no exclusion parameters. In Cytoscape, a first “fraction layout” using the number of scan per fraction was generated to map the relative distribution of each node as pie chart (Fig S2A, in Supporting Information). A second fraction layout using the sum of the precursor ion intensity per fraction was generated using advanced output options of GNPS workflow (Fig. S2B, in Supporting Information). No significant differences were observed between both approaches. The fraction layout showed that the MN annotated as 4-deoxyphorbol esters was over-represented in the most active fractions F7 and F9. The cluster of jatrophane esters of group C was also specific to F9 indicating that 4-deoxyphorbol esters and jatrophane esters of group C could be related to the potent anti-CHIKV activity of F9.

### Semi-preparative SFC-based Purification of Molecules Detected by Molecular Networking

Based on the above MS^2^MN analysis, fraction F9 was selected for further purification of 4-deoxyphorbol esters and jatrophane esters of group C using a carbon-graphite semi-preparative column (Figure 5A). The UV chromatogram at 230 nm was used as proxy (Figures 5B) during the transposition from the analytical to the semi-preparative scales. The conditions of separation were optimized to achieve an adequate collection profile (Figure 5C). The purification of F9 (200 mg) was achieved in 25 runs using a fully automatized batch mode (t ≈ 26 h). A total of 1.5 L of ethanol was used for this purification step. For comparison, the LC purification procedure for the standards was extremely time-consuming (three weeks), and consisted of at least 4 steps in normal phase and reverse phase LC to obtain pure compounds. During this last procedure, the solvent consumption was of approximately 35 L of ethyl acetate, 30 L of methanol, 15 L of heptane, 10 L of acetone, 15 L of acetonitrile, and 10 L of water. In addition, the various solvent evaporation steps were time and energy consuming, especially when large amount of water had to be evaporated. This clearly illustrates the efficient of SFC for the purification of natural products from a complex extract (1 day vs 15 days), while avoiding about 100 L of toxic solvent.

**Figure 5.**
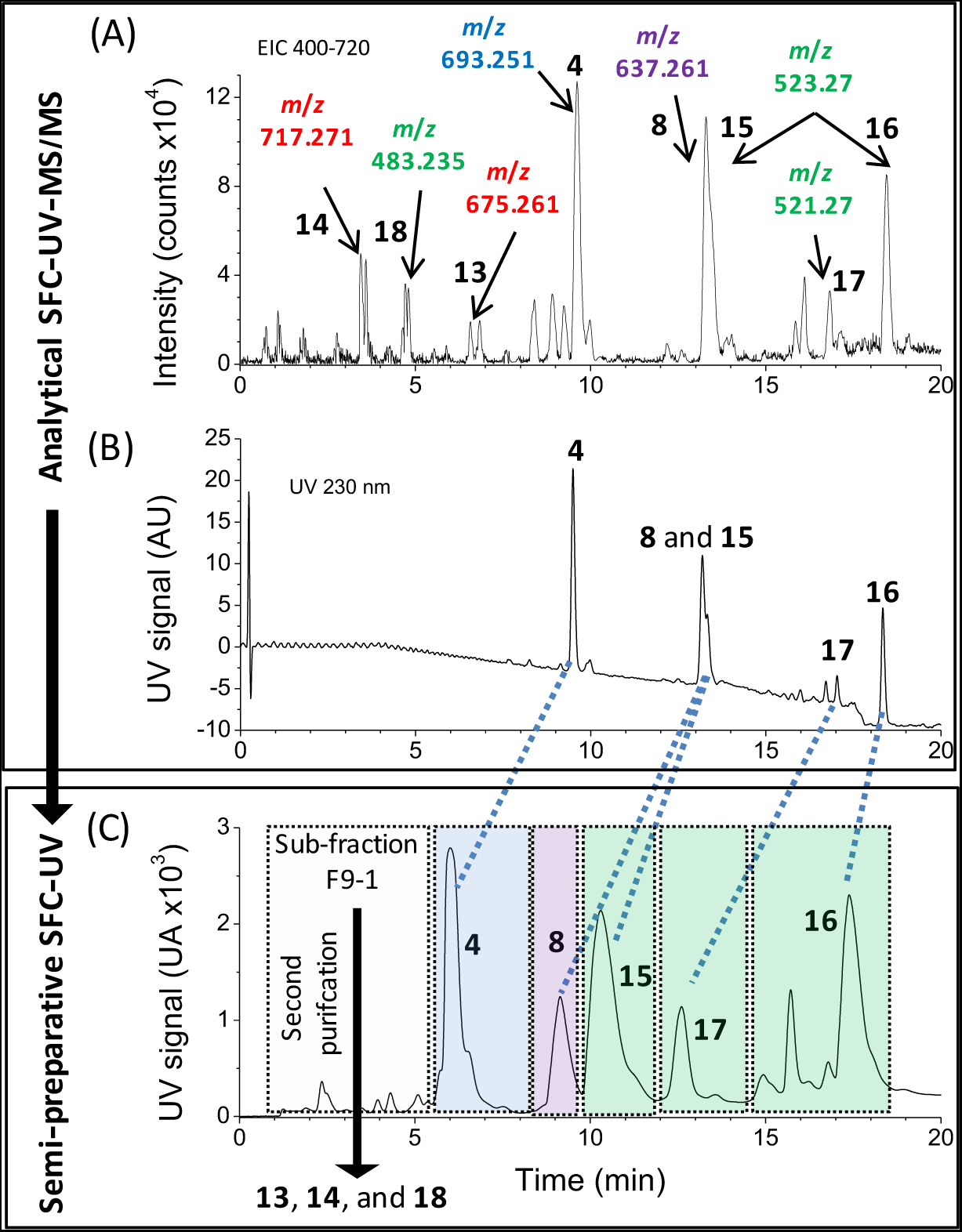
MS-targeted purification procedure by SFC. (A) Extracted ion chromatogram (EIC) of Fraction 9 for all ions between *m*/*z* 400 and *m*/*z* 720. The *m/z* of the targeted and isolated compounds based are indicated. B) The corresponding UV chromatogram at 230 nm. (C) Semi-preparative SFC-UV chromatogram of fraction F9 at 230 nm.

The major peaks at *t*_R_ 9.5 and 13.2 min correspond to compounds **13** (*m*/*z* 693.2510, [C_35_H_42_O_13_+Na]^+^, Δ*m/z* = 0.5 ppm), and **8** (*m/z* 637.2610, [C_33_H_42_O_11_+Na]^+^, Δ*m/z* = 2.3 ppm), respectively The others chromatographic peaks collected at *t*_R_ 18.3, 13.2, and 16.9 min afforded the 4-deoxyphorbol ester **16** (*m/z* 523.2675, [C_29_H_40_O7+Na]^+^, Δ*m/z* = 0.7 ppm) and its epimer **15(** *m/z* 523.2677, [C_29_H_40_O_7_+Na]^+^, *m/z* = 1.0 ppm), and compound **17** (*m/z* 521.2510, [C_29_H_38_O_6_+Na]^+^, Δ*m/z* = 1.0 ppm), respectively. Sub-fraction F9-1 (Figure 5C) was collected based on detection of ions belonging to the cluster of jatrophane of group C. It was further purified using a second semi-preparative SFC leading to the isolation of compound **13** at *m/z* 675.2611 (rt = 6.8 min, [C_32_H_44_O_14_+Na]^+^, Δ*m/z* = 2.6 ppm), compound **14** at *m/z* 717.2711 (rt = 3.8 min, [C_34_H_46_O_15_+Na]^+^, Δ*m/z* = 3.3 ppm), and compound **18** at *m/z* 483.2352 (rt = 4.9 min, [C_26_H_38_O_6_,+Na]^+^, Δ*m/z* = 1.4 ppm). It should be noted that the MS signal of the ion at *m/z* 483.2352 is about 1000 times lower than the signal of the major peak at *m/z* 693.2510 suggesting that the method is efficient for the isolation of abundant and minor secondary metabolites.

### Structural Elucidation of Diterpene Esters Isolated by Semiprep-SFC

Although MS/MS from molecular networks shows the type of modified forms of the diterpenes are present, it cannot provide a full structure. Therefore, the structural elucidation of compounds **13-18** (Figure 3) was achieved using a combination of NMR spectroscopy, ESI-HRMS/MS, and X-ray crystallography (Tables S1-S3, Figures S3 in Supporting Information).

Compounds **13** and **14** are heptaesters 2,3,5,7,8,9,15-heptahydroxylated-oxojatropha-6(17),11*E*-diene, closely related to euphodendroin and amygdaloidin.^34,35^ In their ^1^H NMR spectra, important signal broadness resulting from a slower conformational exchange rate was observed. This phenomenon, caused by the hindering effects of ester groups,36,37 was solved by increasing the analysis temperature as described in SI.

From the analysis of ESI-HRMS, ^1^H and ^13^C NMR data, it was deduced that compounds **15**-**18** were 4-deoxyphorbol esters. Compound **15** was identified through comparison of its ^1^H-NMR and HRESIMS data with those previously reported.^29^ Additional 13C NMR data are provided in Table S2 (Supporting Information). Compound **17** is a 4,20-dideoxyphorbol ester possessing a C-20 aldehyde group. The tendency of phorbol esters to easily oxidize and produce an *α*,*β*-unsaturated carbonyl was previously reported by Schmidt and Hecker.^38^ Thus, **17** could be a direct oxidation product of **15**.

Remarkably, our results showed that MS^2^MN could distinguish MS/MS spectra of macrocylic diterpene esters from phorbol esters. Thus, by overcoming this limitation,^32,33^ MS^2^MN have to be considered as a powerful tool for MS/MS-based profiling of diterpene esters from *Euphorbia*(ceae).

### Evaluation of the Antiviral Activity of the Isolated 4-Deoxyphorbol Esters

Compounds **13**-**18** were evaluated for their anti-CHIKV potential by using a chikungunya virus-cell-based assay, whereas the four 4-deoxyphorbol esters (**15**-**18**) were evaluated for antiviral activity using recombinants HIV-1 (NL4.3-Ren) in MT-2 cells (Table S4, in Supporting Information). Due to the low amount of isolated, compounds **13** and **14** were not evaluated for anti-HIV activity. Their activity was compared to the 12-*O*-tetradecanoylphorbol-13-acetate (TPA), a phorbol ester known for his potent tumor-promoter activity, and prostratin (12-deoxyphorbol-13-acetate),39 a promising non-tumor-promoter antiviral diterpene ester, which is currently under clinical trial evaluation in the US.^40^

Considering the 4-deoxyphorbol esters series, compound **18** was found to be the most potent and selective inhibitor of CHIKV replication (EC_50_ around 440 nM and a selectivity index (SI) > 172). Structure–activity relationships (SARs) were consistent with those already described for the anti-CHIKV activity of phorbol esters,^41^ such as requirement of an *α*-configuration of H-4, and the deleterious effect of a carbonyl group at C-20. In addition, results reported herein showed that the replacement of a tiglyl group by an acetyl group at C-12 greatly increases the selectivity for anti-CHIKV activity (**15** vs **18)**. Compound **15** displayed selective inhibition of HIV-1 with an EC_50_ of 13 nM and SI > 3900. Compound **18** also exhibited a potent and selective antiviral activity with EC_50_ of 54 nM and a SI > 900. Both compounds showed more potency than prostratin (IC_50_ = 226 nM, SI > 221). The other 4-deoxyphorbol esters were significantly less potent (low µM range). The C-4 *β*-configuration looks crucial (**15** *vs* **16**) for an antiviral potency, while an oxidation at C-20 (**15** *vs* **17**) proved to be deleterious. The substitution of a tiglyl group by an acetyl group at C-12 (**15** *vs* **18**) did not modify the activity. Compounds **15** and **18** differ from TPA by the presence of C-12 and C-13 short acyl side chains and by absence of a C-4 hydroxy group. Previous evaluation of tigliane-type diterpenoids for tumor-promoting activity indicated that long C-12 and C-13 acyl side chains were key features for tumor-promoting activity.42,43 Thus, as it is the case for prostratin, compounds **15** and **18** could be devoid of any tumor-promoting activity, making these compounds very attractive for further biological investigation. These results highlight that *Euphorbia* plants are a remarkable source of potent antiviral diterpene esters.^27, 41, 44–46^

In the present study, an SFC-MS/MS based workflow was developed to analyze the diterpene esters content of *E. semiperfoliata*. Data were analyzed through MS^2^MN, leading to (i) the dereplication of known jatrophane esters, (ii) the efficient annotation of analogues, and (iii) the identification of new diterpene esters that were potentially involved in the bioactivity found for the extract. A MS-targeted SFC-purification was carried out leading to the isolation and identification of two new jatrophane esters (**13** and **14**), and one known (**15**), along with three new 4-deoxyphorbol esters (**16-18**). Compounds **15** and **18** exhibited potent and selective antiviral activities against HIV-1 replication, and at a lesser extent against CHIKV replication.

The discovery of new bioactive natural products can thus be accelerated using a SFC-MS/MS-based workflow, while decreasing the chemical risk and reducing the environmental impact of the whole process. Moreover, the present study brings a unique insight on MS^2^MN annotation, by investigating the fragmentation pattern responsible of the clustering of MS/MS spectra in the molecular networks. We demonstrated that MS^2^MN is an efficient tool to explore the molecular content of bioactive fractions, allowing annotation, dereplication of analogues, and prioritization of fractions to be purified. From a methodological perspective, further developments, such as upstream extraction/fractionation using supercritical fluids, would permit a complete eco-friendly workflow, raising metabolomics and natural product chemistry to a Green Era.

## EXPERIMENTAL SECTION

### Vegetal Material, Extraction, and Fractionation

The whole plant of *E. semiperfoliata Viv. (= synonym Euphorbiq qmygdqloides ssp semiperfoliqtq A. R. Sm)* was collected by L.-F. N. in November 2011 near Bocca di Vergio, at an altitude of 1200 m, in the Niolu region of Corsica (France) (GPS coordinates: 42°17′26.999′′ N, 8°54′2.894′ E) and identified by L.-F. N. and Marie-José Battesti. A voucher specimen (LF-023) was deposited at the Herbarium of the University of Corsica. The whole plant of *E. semiperfoliata* was air-dried, ground (dry weight, 370 g), and extracted three times with 5 L of ethyl acetate (EtOAc) at 40 °C and 100 bar using a Zippertex static high-pressure, high-temperature extractor. The EtOAc extract was concentrated under vacuum at 40°C to yield 24.7 g of residue (ρ = 6.7% w/w). The residue was mixed with Celite and subjected to flash LC on a silica column (Versapak, 80 × 150 mm, 20−45 µm, 385 g, acetone−n-heptane 0 to 100% in 60 min, 100 mL/min), leading to ten fractions (F1-F10).

## General Experimental Procedures

### SFC-MS

Supercritical fluid chromatography (SFC) analyses were performed using a 1260 Infinity Analytical SFC system from Agilent Technologies (Waldbronn, Germany) equipped with an Aurora A5 Fusion module, generating supercritical CO_2_ from gaseous CO_2_. A back-pressure regulator (BPR) was used to fix the pressure at 150 bar after the column. Samples were kept at 4°C and 1 µL was injected. SFC was coupled to a diode array detector and to a quadrupole time-of-flight mass spectrometer (Agilent 6540 Q-TOF, Agilent Technologies) equipped with an electrospray ionization source operating in positive mode. Source conditions are provided in Supporting Information. A blank (pure ethanol) was injected after each sample analysis.

A first set of columns was purchased from Agilent Technologies (Massy, France) including ZORBAX Rx-SIL RRHT (Si, 100 × 2.1 mm, 1.8 µm particles), ZORBAX SB-CN RRHT (CN, 100 × 2.1 mm, 1.8 µm particles), Pursuit pentafluorophenyl (PFP, 150 x 2.0 mm, 3.0 µm particles), Eclipse ZORBAX Eclipse Plus C_18_ RRHT (C_18_, 150 × 2.1 mm, 1.8 µm particles) and Pursuit XRs Diphenyl (DPH, 250 × 2.0 mm, 3 µm particles). A second set of column was provided by Waters (Guyancourt, France) including ACQUITY UPC² BEH 2-EP (2-ethylpyridine) (2-EP, 100 × 2.1 mm, 1.7 µm particles), ACQUITY UPC² Torus 1-AA (1-Aminoanthracene), ACQUITY UPC² Torus 2-PIC (2-Picolylamine) and ACQUITY UPC² Torus DEA (Diethylamine) (150 × 2.1 mm, 1.7 µm particles). Finally, a Discovery^®^ Zr-Carbon column (Zr-Carbon, 150 × 2.1 mm, 3 µm particles) was purchased from Supelco Saint-Quentin Fallavier, France) and Hypercarb^®^ column (Carbon, 100 × 2.1 mm, 3 µm particles) from Thermo Scientific (Courtaboeuf, France). The column oven was maintained at 30°C. The porous graphitic carbon column (Hypercarb^®^) was finally selected after column screening. The elution gradient was optimized using CO_2_ (solven and ethanol with 0.1 % formic acid (solvent B). The optimal gradient was: 0–3 min at 3 % solvent B, 3–13 min linear gradient from 3 to 10 % solvent B, 13–17 min from 10 to 20 % solvent B, 17–20 min isocratic plateau at 20 % solvent B, 21-23 min re-equilibration at 3 %, at a constant flow rate of 1.5 mL/min.

### SFC-MS/MS method

Untargeted MS/MS analyses were performed using an optimized data-dependent acquisition (DDA) mode consisting of a full MS scan from *m/z* 100 to *m/z* 1000 (scan time: 100 ms), followed by DDA of MS/MS spectra of the four most intense ions (Top-4, 250 ms each) from *m/z* 300 to *m/z* 800, with a minimum intensity of 2000 counts. No exclusion rule was used to use the number of scan as pseudo-quantitative proxy in MN2MS. Collision energy was fixed at 30 eV according to the optimized collision-induced dissociation (CID) conditions (Figure S1, in Supporting Information).

#### Molecular networks (MS^2^MN)

Molecular networks (MNs) were created using the Data Analysis workflow on GNPS platform (http://gnps.ucsd.edu).^19^ The parameters used are detailed in Supporting Information. For the clustering of consensus MS/MS spectra, parameters were set for DE with a parent mass tolerance of 0.01 Da (allowing an error of 25 ppm at *m*/*z* 400 and 12 ppm at *m*/*z* 800 for the precursor ion), a product ion tolerance of 0.0075 Da allowing a maximum error of 25 ppm at *m*/*z* 300 and 12 ppm at *m*/*z* 600). The fragment ions below 10 counts were removed from MS/MS spectra. MNs networks were generated using 12 minimum matched peaks and a cosine score of 0.7. MNs were visualized using Cytoscape 3.4.0 software. ^47^ A force-directed layout modulated by cosine score factor was used for data visualizations. Semi-quantitative estimation was provided by representing the size of the node by the sum of precursor ion intensities and by mapping the number of scan for each fraction as pie-chart diagram. MS/MS data were deposited to MassIVE Public GNPS dataset (http://gnps.ucsd.edu, MSV000079856). Molecular networks were created using the online workflow at GNPS. The data were filtered by removing all MS/MS peaks within +/-17 Da of the precursor m/z. MS/MS spectra were window filtered by choosing only the top 6 peaks in the +/- 50Da window throughout the spectrum, and every fragment ions below 10 count in the MS/MS spectra were removed. The data was then clustered with MS-Cluster with a parent mass tolerance of 0.01 Da and a MS/MS fragment ion tolerance of 0.0075 Da to create consensus spectra. A network was then created where edges were filtered to have a cosine score above 0.7 and more than 12 matched peaks. Further edges between two nodes were kept in the network if and only if each of the nodes appeared in each other's respective top 10 most similar nodes. The spectra in the network were then searched against GNPS's spectral libraries. The library spectra were filtered in the same manner as the input data. All matches kept between network spectra and library spectra were required to have a score above 0.7 and at least 6 matchedpeaks. Defaultparameters: http://gnps.ucsd.edu/ProteoSAFe/status.jsp?task=61a9770e433043a6a88ab125a2cffb7c and optimized parameters: http://gnps.ucsd.edu/ProteoSAFe/status.jsp?task=90e3d8b4fe7545fd93c16e950d1686f7

### Relative Distribution Mapping in Molecular Networks

Using the advanced output option of Data Analysis GNPS workflow, “Create cluster bucket”, a fraction layout was created. The output bucket table was imported into Cytoscape. In Cytoscape, the size of the node was modulated by the total sum of precursor ion intensities, and the piechart diagram represent the sum of precursor ion intensities in each fraction (Figure S7) or the number of scan (Figure S8).

### Visualisation of Molecular Networks in Cytoscape

Data were visualized using Cytoscape 3.4.0 software.^47^ A force-directed layout (Allegro Weak Clustering, http://allegroviva.com/allegrolayout2) modulated by cosine score factor was used for data visualizations. In Cytoscape, molecular networks with less than 3 components were not considered, in order to focus one main spectral families. Couple of nodes corresponding to isotopologues ^13^C were manually discarded. Allegro Layout Weak clustering parameters: Algorithm tuning: scale (75%) and gravity (circular, 50%). Edge weighting parameters were set as default except weight transform (Cosine, inverse log: x =>-Log10(x). Prevent nodes from overlapping.

### Semi-preparative-SFC and Purification Procedure

A semi-preparative SFC (semiprep-SFC) system from Waters^®^ (Milford, Massachusetts, USA) was used for compound isolation. It consists of a SFC fluid delivery module, an autosampler with a 48-vial plate, a SFC analytical-2-prep oven with a 10-column selection valve, a SFC BPR SuperPure Discovery Series from Thar and a cooling bath of type Neslab RTE7 controlled by a Digital One thermoregulator from Thermo Scientific. The system is coupled to a PDA 2998 and a DEDL 2424 from Waters. The autosampler was equipped with a 100 µL injection loop. A porous graphitic carbon column (Hypercarb, 150 x 10 mm, 5 µm particles was used. An automated six-vessels collection module was employed with a solvent flow set to 3 mL/min. The BPR was fixed at 150 bar, the oven temperature at 40°C, and ethanol was added as co-solvent. The instrument was controlled by Superchrom and data were processed using Chromscope (Thar, Pittsburgh, PA, USA).

### Structural elucidation

Optical rotations were measured using a JASCO P1010 polarimeter at 25 °C. The UV spectra were recorded using a Perkin-Elmer Lambda 5 spectrophotometer. 2D NMR spectra were recorded using a Bruker 500 MHz instrument (Avance 500), and using a Bruker 300 MHz instrument (Avance 300) for ^13^C NMR spectra. CDCl_3_ was used as solvent. Crystallographic data were collected using a Rigaku diffractometer constituted by a MM007 HF copper rotating-anode generator, equipped with Osmic confocal optics, and a Rapid II curved Image Plate. Crystallographic data for **16** have been deposited to the Cambridge Crystallographic under the reference 1491995.

### Compound properties

*(2R,3R,4S,5R,7S,8R,9S,13R,15R)-3,5,7,9,14,15-Hexaacetoxy-8-hydroxy-14-oxojatropha-6(17),11E-diene* (**13**, JE-C1): Amorphous powder; [*∝*]D^25^ −95 (*c* 0.1, MeOH). For ^1^H and ^13^C NMR spectroscopic data, see Table S1; HRESIMS *m/z* 675.2613 [M + Na]+ (calcd for C_32_H_44_O_14_, ∆*m/z* theoritical = + 1.4 ppm); (HRESIMS/MS deposit to GNPS spectral library, ref: CCMSLIB00000531511); See S12-S19 in Supporting Information.

*(2R,3R,4S,5R,7S,8R,9S,13R,15R)-3,5,7,8,9,14,15-Heptaacetoxy-14-oxojatropha-6(17),11E-diene* (**14**, JE-C2): Amorphous powder; [*∝*]D^25^ −130 (*c* 0.5, MeOH). For ^1^H and ^13^C NMR spectroscopic data, see Table S1; HRESIMS *m/z* 717.2733 [M + Na]+ (calcd for C_32_H_45_O_13_, ∆*m/z* theoritical = + 0.1 ppm); (HRESIMS/MS deposit to GNPS spectral library, ref: CCMSLIB00000531512); See 20-S29 in Supporting Information.

*4∝-deoxyphorbol 12-tiglate-13-isobutyrate* (**15**, 4-dPE A): Amorphous powder; [*∝*]D^25^ + 15 (*c* 0.1, MeOH). For ^1^H and ^13^C NMR spectroscopic data, see Table S2; ESI-HRMS *m/z* 523.2674 [M + Na]^+^ (calcd for C_29_H_40_O_7_, ∆*m/z* theoritical = −0.9 ppm); (ESI-HRMS/MS deposit to GNPS spectral library, ref: CCMSLIB00000531515); See S30-S37 in Supporting Information.

*4∝-deoxyphorbol 12-tiglate-13-isobutyrate* (**16**, 4∝-dPE A): Colorless crystal; [*∝*]D^25^ + 30 (*c* 0.1, MeOH). For ^1^H and ^13^C NMR spectroscopic data, see Table S2; ESI-HRMS *m/z* 523.2675 [M + Na]^+^ (calcd for C_29_H_40_O_7_, ∆*m/z* theoritical = −0.7 ppm); (ESI-HRMS/MS deposit to GNPS spectral library, ref: CCMSLIB00000531518); See S38-S45 in Supporting Information.

4∝,20-dideoxyphorbol12-tiglate-13-isobutyrate(**17**,4,20-ddPEA): Amorphous powder; [∝]D^25^ + 20 (c 0.1, MeOH). For ^1^H and ^13^C NMR spectroscopic data, see Table S3; ESI-HRMS m/z 521.2517 [M + Na]^+^ (calcd for C_29_H_38_O_7_, ∆m/z theoritical = −1.2 ppm); (ESI-HRMS/MS deposit to GNPS spectral library, ref: CCMSLIB00000531514); See S46-S53 in Supporting Information.

4∝-deoxyphorbol 12-acetate-13-isobutyrate (**18**, 4-dPE B): [∝]D^25^ + 40 (c 0.1, MeOH). For ^1^H and ^13^C NMR spectroscopic data, see Table S3; ESI-HRMS m/z 483.2348 [M + Na]+ (calcd for C_26_H_36_O_7_, ∆m/z theoritical = + 1.3 ppm); (ESI-HRMS/MS deposit to GNPS spectral library, ref: CCMSLIB00000531516); See S54-S61 in Supporting Information.

### Antiviral Assays

The anti-Chikungunya virus (CHIKV) bioassays were achieved according to the protocols previously described.41 Anti-HIV bioassays were performed as detailed in the Supporting Information.^48^

## ASSOCIATED CONTENT

### Supporting Information

The Supporting Information is available free of charge on the ACS Publications website at DOI: XXXX. LC-ESI-HRMS/MS data, and MS/MS spectra of compounds are available on GNPS spectral library (http://gnps.ucsd.edu). 1D and 2DNMR spectra are shown in Supporting Information. Crystallographic information file (CIF) for compound 16 is provided. This material is available free of charge via theInternet at http://pubs.acs.org.

## ACKNOWLEDGMENT

This work has benefited from an “Investissement d’Avenir” grant managed by Agence Nationale de la Recherche (CEBA, ANR-10-LABX-25-01)). Financial support was provided to LFN and PD by the National Institutes of Health (NIH) grant GM097509. JA was supported by The SPANISH AIDS Research Network RD12/0017/0015 that is included in Acción Estratégica en Salud, Plan Nacional de Investigación Científica, Desarrollo e Innovación Tecnológica 2008-2011, Instituto de Salud Carlos III, Fondos FEDER and Instituto de Salud Carlos III-FIS: PI12/00506. EDLT was supported by a fellowship of the Instituto de Salud Carlos III (ISCIII-FIS N°. FII4CIII / 00014). Authors would like to thank Odile Thoison (CNRS-ICSN) for her precious help with semipreperative-SFC experiments.

